# Improving the Hodgkin-Huxley Models of Ionic Conductance and Action Potential Generation

**DOI:** 10.64898/2026.08.04.742717

**Authors:** Moussa Djioua

**Author notes:** Contributing authors.

## Abstract

This study presents improvements to the Hodgkin-Huxley (HH) models of ionic conductance and action potential generation. Sodium and potassium conductances are expressed by a single analytical formula describing the impulse response of a convolution of exponential distributions within a short-memory integration space. Treating transmembrane ion transit duration as a random variable, conductance profiles are interpreted as realizations of the probability density functions governing ionic movements. Applying the central limit theorem, the lognormal distribution emerges as the asymptotic profile of ionic conductances, constituting a fundamental primitive for such biosignals. A temporal state-transition paradigm describes the action potential waveform through four successive membrane potential transitions. Applied to electrophysiological recordings from lamprey reticulospinal neurons, this framework enables indirect estimation of key physiological quantities, including depolarization threshold, Nernst potentials, and net ion fluxes across the membrane. These advances open new perspectives for parameter estimation from experimental data and neuronal network simulation.

## 1 Introduction

In voltage-clamp experiments, Hodgkin and Huxley modeled the ionic conductance of the giant axon of *Loligo* using functions expressing the activation and inhibition probabilities of ion channel gates, reflecting the conditions for ionic translocation across the membrane. A system of first-order differential equations was then proposed to reproduce the action potential (AP) waveforms arising from brief membrane depolarization [1–5]. Improving this model is motivated by the need to (1) better understand the biological mechanisms underlying the membrane’s response to depolarization, (2) analyze biosignals recorded during motor or other tasks, (3) model neuronal dysfunction resulting from ionic imbalances, and (4) study neuronal behavior within networks, with a view to constructing bio-inspired systems that replicate human tasks.

The present study summarizes research conducted to improve the original HH models on two fronts. First, regarding ionic conductance, a re-examination of the original HH equations and data led to reformulating the model using a single analytical function, validated through curve fitting. Second, regarding the AP waveform, a state-transition paradigm was used to describe it with a single analytical equation composed of four successive transitions corresponding, respectively, to the presynaptic (PSP), depolarization (DEP), repolarization (RPP), and return-to-rest (RRP) phases of the membrane potential. This model was applied to fit AP trains recorded from lamprey reticulospinal neurons, demonstrating its capacity to extract the structure of the underlying biological system and to estimate key biophysical quantities, including Nernst potentials and the depolarization threshold.

A discussion addresses the stochastic nature of the processes involved at multiple levels in the deterministic electrical responses of biological systems, with particular focus on the neuron.

## 2 Results

### 2.1 The Hodgkin-Huxley Model

From Hodgkin and Huxley’s voltage-clamp experiments [1–5], ionic conductance of the neuronal membrane has been modeled by a system of coupled first-order equations yielding analytical expressions that resemble probability densities, faithfully reproducing the characteristic profiles of sodium conductance *g*_*Na*_, potassium conductance *g*_*K*_, and the AP. Since then, several mathematical models have been proposed to describe the AP waveform [7, 8]. The HH equations are here re-examined from three complementary perspectives: the stochastic nature of biological system functioning, the central limit theorem (CLT)— which describes how deterministic behavior emerges from stochastic processes — and the agonist-antagonist synergy inherent to any system where every direct action elicits an opposing reaction. Indeed, the deterministic AP waveform arises from ionic movements that are stochastic at the molecular level, while the presence of ion pumps and the opposing flows of sodium and potassium ions exemplify such antagonistic couplings.

We first ask whether the ionic conductance profile is stereotypical and independent of ion type. The membrane’s ionic permeability constitutes a biological system comprising a large number of parallel ion channels, each formed by a series of gates representing successive conditions of ionic passage. Since ion transit through a channel is a causal, ergodic stochastic process, the CLT implies that the deterministic time profile of conductance is the direct realization of the probability density function governing the duration of ionic translocation across the membrane. In the sense of the law of large numbers, what is measured at the macroscopic scale is simply the average deterministic behavior of an underlying microscopic stochastic process.

This interpretation is supported by the HH model itself: sodium conductance is expressed as a product of gate probabilities, yielding a profile analogous to the Erlang function — a probability density arising from the convolution of *n* exponential processes, combining an activation term (power-law rise) with an inhibition term (decaying exponential). Bell-shaped conductance curves observed in AP simulations [4] further confirm that both conductances can be modeled as probability density functions.

#### 2.1.1 Ionic Conductance Model

The models proposed by Hodgkin and Huxley to describe the time courses of the sodium *g*_*Na*_ and potassium *g*_*K*_ conductances of the neuronal membrane are derived from a system of coupled first-order differential equations, whose solutions are given by the following analytical expressions (Equations 11 and 19 in [4], pp. 508 and 513, respectively).

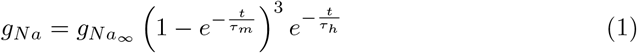

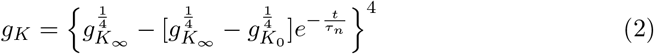

Since, at rest, the conductance *gK*_0_ = 0, we obtain a sigmoid-like curve that can be rewritten as follows:

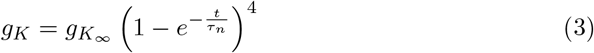

Fitting the experimental data yielded exponents of 3 and 4 for the sigmoid-like activation functions of sodium and potassium conductances, respectively. Cole and Moore [6] later proposed setting the potassium exponent *n* to 25 and introduced an onset time *t*_0_ relative to depolarization. Furthermore, Tables 1 and 2 of [4] reveal highly proportional relationships between the time constants: *τ*_*h*_ ≈ 3*τ*_*m*_ for sodium and *τ*_*n*_ ≈ 7*τ*_*m*_ for potassium, where *τ*_*m*_ is the activation time constant of the sodium channel gates. The potassium activation time constant is thus approximately 7 times longer than that of sodium. These observations motivate a unified analytical representation for both conductances by a density profile described by:

**Table 1:** Summary of the Mean Values of Selected Markers Characterizing the Four State Transitions.

| phase | $t_0$<sup>1</sup>(ms) | $t_r$ | $t_x$ | $t_{\max}$ | $t_i$ | $T$ | $E_n$ (mV) | $Q$ (pmol) |
| --- | --- | --- | --- | --- | --- | --- | --- | --- |
| Presynaptic (PSP) | 0 | 10.0 | 13.9 | 17.2 | 10.0 | 21.2 | 39.5 | 0.20 |
| Depolarization (DEP) | 14.6 | 15.5 | 15.9 | 16.4 | 0.9 | 3.6 | 65.4 | 0.92 |
| Repolarization (RPP) | 15.4 | 16.0 | 16.5 | 17.4 | 0.6 | 11.5 | -89.8 | -1.63 |
| Return to rest (RRP) | 16.8 | 17.0 | 17.1 | 17.7 | 0.2 | 46.6 | 76.7 | 0.47 |

<sup>1</sup>The onset time $t_0$ of the PSP phase, expressed in (ms), serves as the temporal reference for all markers except $T_m$ and $t_i$ . Markers associated with potassium ions are assigned a negative sign to indicate the direction of the corresponding ionic current

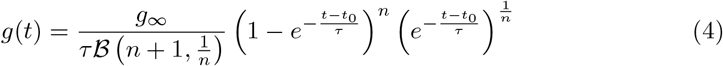

where *τ* is the time constant of the exponential law governing ionic transit, *t*_0_ the depolarization onset time, 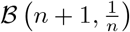 Beta constant, and *g*_∞_ a constant proportional to the total number of ions crossing the membrane. Setting *n* = 3 yields equation (1); setting *n* = 25 with *τ*_*n*_ ≫ *τ*_*m*_ yields equation (3), since *e*^−*t/*25*τ*^ ≈ 1.

Sigworth and Neher [9] showed that transmembrane ion transport produces a discrete current that randomly switches between two states, forming a spike train with exponentially distributed interspike intervals. This establishes membrane permeability as an ergodic, causal stochastic process, so that the conductance measured in voltage-clamp experiments is the direct realization of a probability density — specifically, the density of a sum of i.i.d. exponential random variables. Using the equivalence between stochastic processes and linear systems introduced by Papoulis [10], the conductance of equation (4) is expressed as a convolution product of exponential impulse responses, defined within a short-memory integration space endowed with the Lebesgue measure *µ*(*t*) = 1 − *e*^−*αt*^.

#### 2.1.2 Analysis of the Conductance Model

The analytical expression of the conductance in (4) admits several interpretations.

In the first interpretation, the change of variables 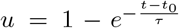 yields a beta-distributed conductance profile, such that:

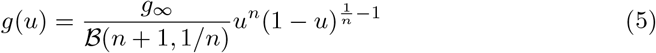

This probability density describes a positively skewed profile with a bounded, normalized variable on [0, 1). The change of variables maps the time domain [0, ∞) onto a normalized space, where *u* represents the relative advancement of the conductance activation process, evolving logarithmically as *t* = − *τ* ln(1 − *u*). Since *n* + 1 *>* 1*/n* in the HH conductance model, the density reaches its maximum rapidly, yielding a positively skewed profile. By analogy with Papoulis’ approximation of the symmetric beta density to the Gaussian [10], this family of asymmetric beta densities may be regarded as a good approximation to the lognormal distribution.

In a second interpretation, the model expresses the set of conditions for ion channel opening as a convolution of *n* + 1 i.i.d. exponential densities with decay time constant *nτ* , endowed with the saturating Lebesgue measure *µ*(*t*) = 1 − *e*^−*t/τ*^ . This measure reflects the short-memory nature of the integration medium: ion channels remain active only transiently, with throughput decreasing exponentially over time. Two distinct stochastic mechanisms are thus identified — transient channel activation and ionic translocation — both contributing to membrane conductance following depolarization.

Following Papoulis’ equivalence between independent random variables and system impulse responses [10], this kind of convolution corresponds to the sum of *n* + 1 i.i.d. exponential random variables sharing the same time constant *nτ* . The inactivation introduced by HH reflects the transient nature of depolarization-induced channel opening: even under constant voltage, as in voltage-clamp conditions, the channel’s permeability decays over time. This nonlinear, threshold-dependent mechanism is represented by *µ*(*t*), indicating that channel throughput decreases as ions traverse the membrane.

The second mechanism corresponds to the ion channel structure itself, which, according to the HH model, consists of a set of temporal gates whose mean open duration depends on their number. Each gate is modeled as a first-order RC system with time constant *τ* . When interconnected within a network, the effective time constant increases to *nτ* , suggesting a coupled architecture in which each gate is connected to the remaining *n* − 1 gates. The ion channel thus behaves as a diffusive network: passive RC coupling — via either an equivalent parallel capacitance *C*^*′*^ or a series resistance *R*^*′*^ — produces a collective slowing-down effect on ionic transit, such that *τ* ^*′*^ = *R*(*C* + *C*^*′*^) or *τ* ^*′*^ = (*R* + *R*^*′*^)*C*, respectively. The HH conductance equations therefore provide insight into both the structural organization and the collective dynamics of ion channels.

Precisely, through the change of variables on equation (4), yieldings to a beta-distributed conductance profile, by keeping *t*_0_ = 0, the mean conductance is given by: 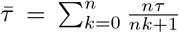. This result indicates that the membrane conductance can be described by a set of n + 1 independent exponential subsystems with distinct time constants 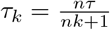 decreasing over convolution level k.

Starting from the assumption that ion channels contain gates opening only when *n* + 1 independent conditions are satisfied simultaneously, the shape of the conductance profile suggests a hierarchical channel structure whose inertia is *n* + 1 times that of an isolated unit. The structure therefore has an incompressible minimum propagation time *nτ* , an additive stochastic processes, to which further conditions proportional to this minimum are added. This points to a highly interconnected, hierarchical organization that can be interpreted as a multiplicative stochastic processes with *n* + 1 variables.

#### 2.1.3 Ion Transit Time as a Random Variable

Ionic conductance expresses the transmembrane permeability of the neuron with respect to various ions. The form of the conductance in equation (4) identifies it as the probability density of a stochastic time variable *T* representing the transit time of an ion across the membrane, such that the instantaneous conductance value gives the number of ions crossing in the same interval.

At the channel exit, the interspike interval — the time between two consecutive ion passages — is random, since the gate structure of the ion channel causes each ion to follow a different pathway. The total transit time, measured from depolarization onset, equals the ion’s intrinsic passage duration plus the cumulative delays introduced by preceding ions.

Transit duration is further governed by the ion’s kinetic energy acquired during depolarization: greater energy accelerates passage but increases random interactions with channel gates. When no gate interaction occurs, the transit time *T*_0_ follows an exponential distribution with mean *τ* , and the total time relative to depolarization is a sum of i.i.d. variables following an Erlang distribution. When the ion interacts with a gate, it loses kinetic energy and its transit time increases by a factor (1 + *ε*_*k*_), where *ε*_*k*_ is a random variable with mean 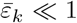 1. After *n* successive interactions, the total transit time becomes the multiplicative product 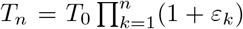 By the central limit theorem, as *n* grows large, *T*_*n*_ converges to a lognormal distribution — the distribution arising from cascades of multiplicative random events.

Let us now relate this proposed multiplicative structure to the additive structure of ion channels suggested by the HH model. Consider the particular case where the *n* variables *ε*_*k*_ have mean value 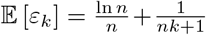, with *n* ≫ 1. Here,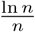 represents the global hierarchical time dilation common to all interaction levels, and 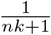 the local time dilation specific to interaction level *k*. The mean transit time *T*_*n*_ then becomes:

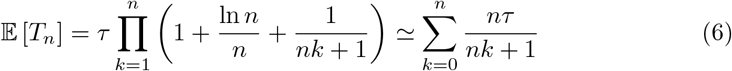

After approximations applied to the multiplicative representation of the stochastic process governing transmembrane ion passage, this particular case indicates that the ionic transit time can be represented by an additive random variable, with variables whose mean is equal to 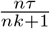 It is precisely this mean value that appears in Equation (4) of the HH ionic conductance model. Thus, this latter may be interpreted as an approximate version of the limiting lognormal profile.

Since the total transit time, measured relative to the depolarization’s onset time, is equal to the sum of its intrinsic duration and the durations associated with the ions that preceded it, it corresponds to the sum of i.i.d. lognormally distributed random variables. Using the Fenton-Wilkinson approximation, this total duration may also be assumed to follow a lognormal distribution.

On the other hand, the fact that the membrane consists of a very large number of ion channels arranged in parallel, together with the causal and ergodic nature of the stochastic processes underlying transmembrane permeability, leads to interpreting the measured conductance as a direct estimate of the density of a lognormal random variable. In other words, the deterministic conductance waveform observed experimentally through ionic current measurements obtained using the voltage-clamp technique corresponds to the probability density function of a random variable associated with the stochastic process governing ionic transmembrane transit during depolarization.

### 2.2 Lognormal Modeling of Ionic Conductance

This modeling approach is further justified by the nature of the voltage-clamp measurement: conductance is recorded via electric current, which represents the rate of ionic transit across the membrane. Since ionic current is fundamentally a rate, we draw on theoretical frameworks developed for velocity profiles of rapid movements, which rely on probability density functions such as the lognormal, beta, and Gaussian functions [14]. In particular, the kinematic theory of rapid movements uses the lognormal function as a fundamental primitive for velocity profile modeling [13]. Ionic conductance, defined as the number of ions crossing the membrane per unit time, is therefore described by the following equation:

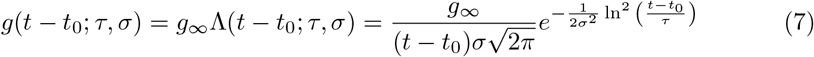

The conductance profile is shaped by the two intrinsic parameters (*τ, σ*) of the lognormal function Λ(*t*), shifted by the onset time *t*_0_ relative to depolarization and scaled by 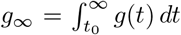 *dt*, which represents the charge transferred per volt — the equivalent transient membrane capacitance, expressed in mF/cm^2^. The parameter *τ* is the geometric mean (equivalently, the arithmetic median) and corresponds to the time at which half of the ions have crossed the membrane. The parameter *σ* is the arithmetic standard deviation and serves as a measure of skewness.

As illustrated in Fig. 4, the conductance duration is subdivided into three phases. The first, *t*_0_, is the delay between depolarization onset and the membrane response. The second, *t*_*i*_ = *τe*^−3*σ*^, is the inertia delay for ion channel opening. The third, *T* = 2*τ* sinh(3*σ*), is the effective conductance duration. The total reaction time is thus *t*_*r*_ = *t*_0_ + *t*_*i*_.

#### 2.2.1 Analysis of the Original HH Data

Figs. 1 and 2 show the fitting results for the sodium and potassium conductance profiles of [4] using the lognormal model of equation (7). Fig. 3 illustrates the dependence of sodium conductance on depolarization voltage, revealing two critical thresholds. Below 25 mV, membrane permeability remains negligible. Between 25 and 75 mV — the normal operating regime — the equivalent transient capacitance *g*∞ increases sigmoidally from approximately 5 to 20 mF/cm^2^, while the temporal markers *τ* , *T* , and *t*_*r*_ decrease exponentially. The onset time *t*_0_ ≈ −0.175 ms remain approximately constant. Beyond 75 mV, *g*_∞_ decreases abruptly while ionic transit accelerates, supporting the hypothesis that high depolarization levels act as an inhibitory mechanism for sodium permeability. For potassium conductance, the recording durations in [4] are insufficient for precise parameter estimation.

**Fig. 1:**
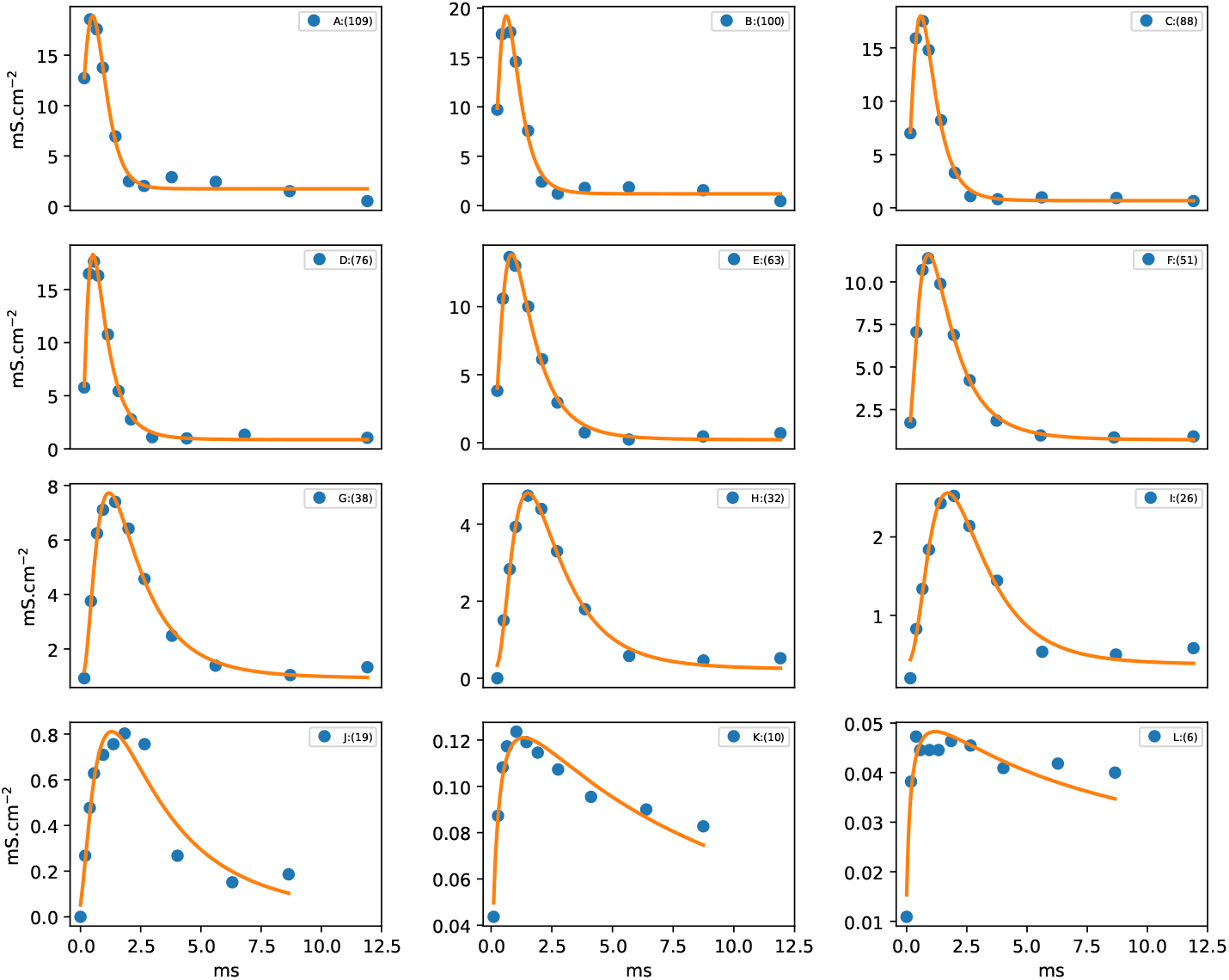
Lognormal fits to sodium conductance recordings obtained at different depolarization levels in the voltage-clamp experiment of Fig. 6 (p. 513) in [4].

**Fig. 2:**
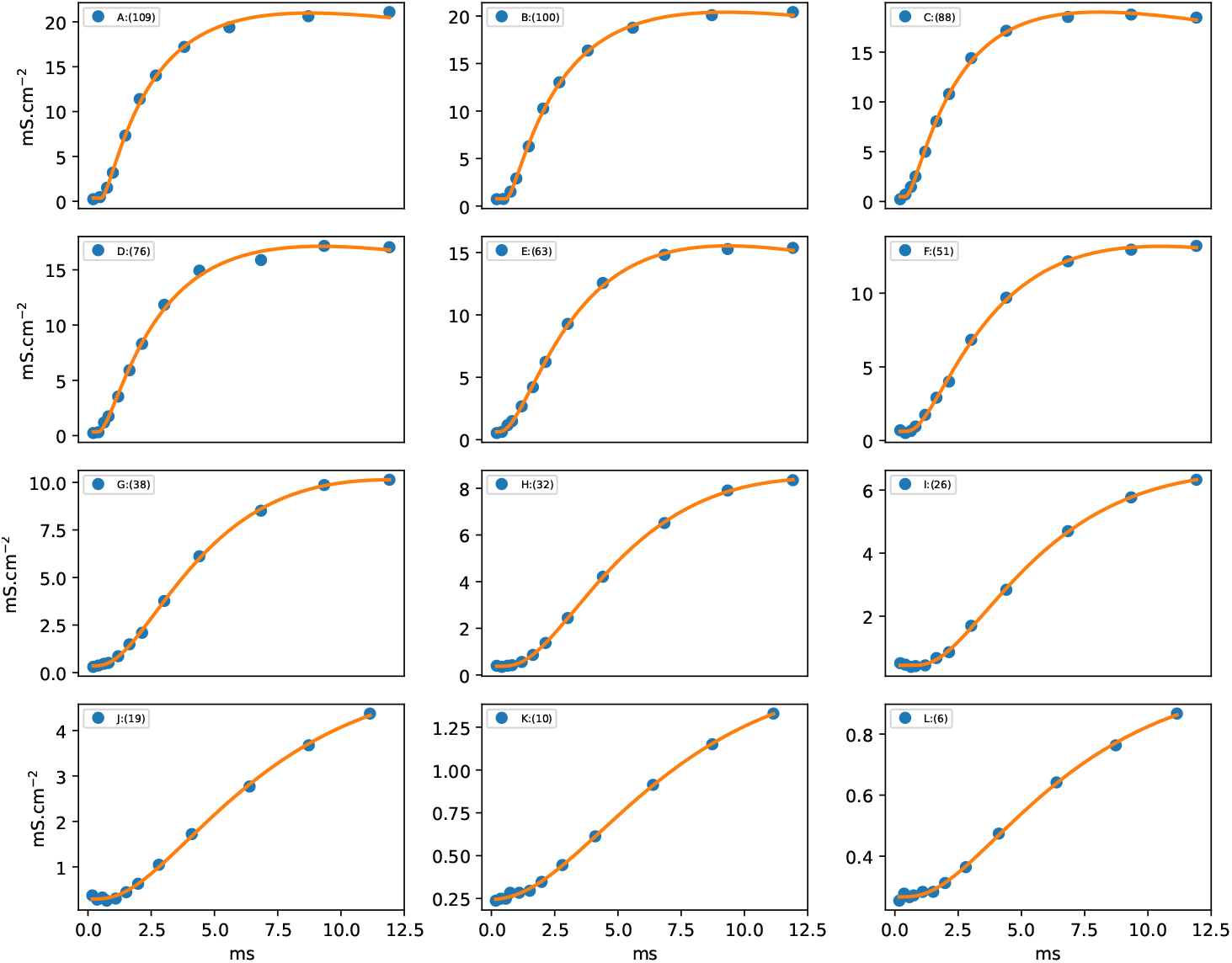
Lognormal fits to potassium conductance recordings obtained at different depolarization levels in the voltage-clamp experiment of Fig. 3 (p. 508) in [4].

**Fig. 3:**
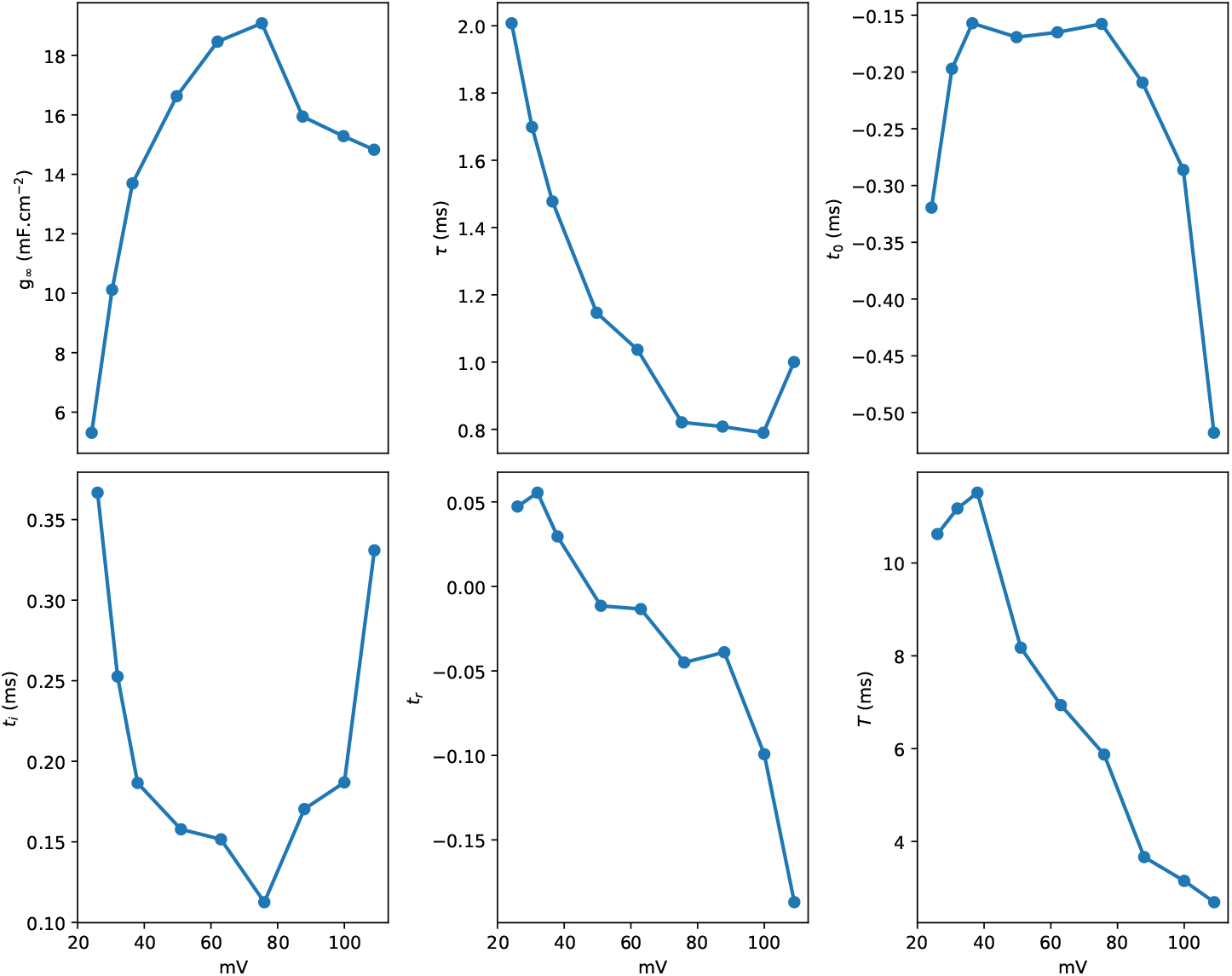
Dependence of selected markers on depolarization voltage, derived from the lognormal sodium conductance model.

**Fig. 4:**
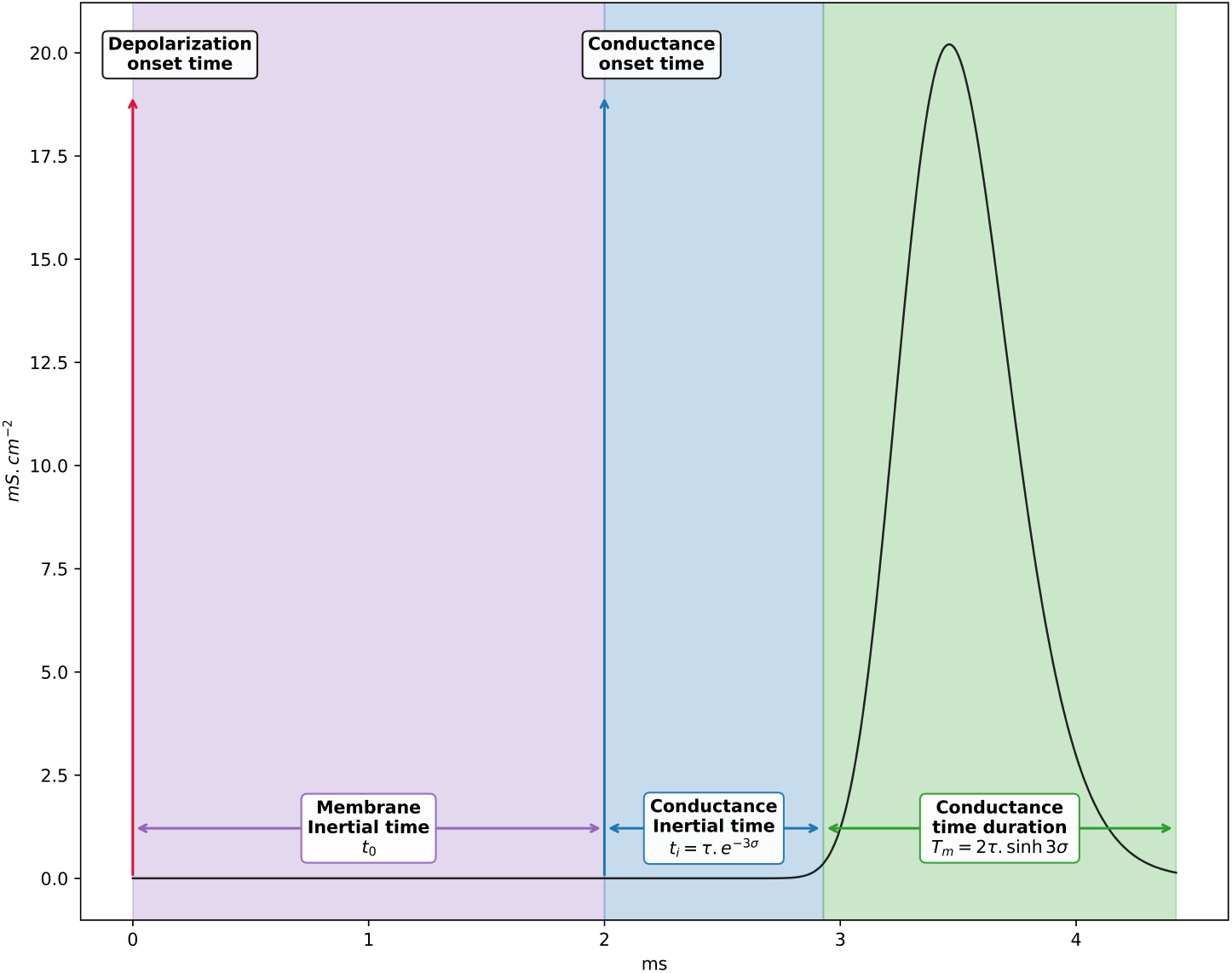
Temporal dilations transforming the membrane response to a step depolarization into a bell-shaped conductance profile.

### 2.3 Action Potential Model

The second component of the HH model describes the AP waveform through the relationship between ionic currents and the charging of the membrane capacitor *C*_*m*_:

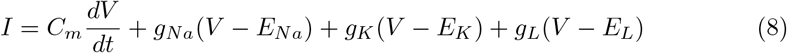

Under voltage-clamp conditions, *V* is held constant and the measured current yields the stereotypical conductance waveforms. Under physiological conditions, the free membrane response to a depolarizing stimulus — modeled as a Dirac impulse — is given by:

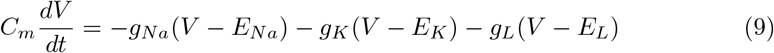

Since conductances are positive, the sodium current *i*_*Na*_ = −*g*_*Na*_(*V* − *E*_*Na*_) *>* 0, as the Nernst potential *E*_*Na*_ = 65 mV constitutes the upper limit of the AP. Conversely, *i*_*K*_ = −*g*_*K*_(*V* − *E*_*K*_) *<* 0, since *E*_*K*_ = −90 mV is its lower limit. With *E*_*L*_ = −65 mV lying between these extremes, the leakage current *i*_*L*_ undergoes two sign reversals at *V* = *E*_*L*_, occurring before depolarization and after repolarization — phases in which leakage is predominant.

The membrane potential *V* is modeled as a sequence of four successive state transitions. The first, the PreSynaptic Phase (PSP), represents the increase in *V* due to an external ionic flux; once *V* exceeds the threshold *V*_*th*_, depolarization is triggered. The second, the Depolarization Phase (DEP), is driven by a rapid influx of sodium ions. When *V* reaches a second threshold, the Recovery Phase (RCP) begins, driven by a prolonged efflux of potassium ions until *V* falls below the resting potential *V*_*rest*_. This phase encompasses both repolarization (REP) and hyperpolarization afterpotential (HAP), here treated as a single state transition dominated by K_+_ efflux. Finally, the Return to Rest Phase (RRP) describes the action of Na/K/ATPase pumps restoring *V* toward *V*_*rest*_ from the hyperpolarized state.

This model assumes that in each phase a single ionic flux is predominant, while others remain approximately constant or absent. The AP profile is thus described as four concatenated transitions with partial overlaps. Letting *V*_*n*_ denote the phase dominated by the conductance of a given ion, and using equations (7) and (9), the current *i*_*n*_ is approximated by:

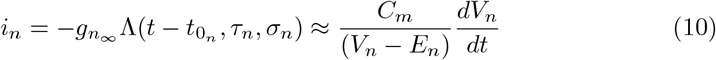

Which gives:

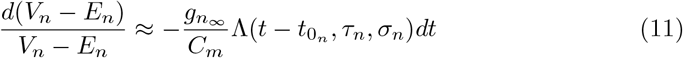

Each state transition driven by a given ionic current occurs at a voltage 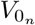 , expressed as:

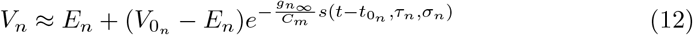

According to equation (11), *g*_*n*∞_ has units of mF/cm^2^ and *C*_*m*_ is in *µ*F/cm^2^, so the ratio *α n* = *g*_*n*∞_ */C* _*m*_ is dimensionless, with a factor of 10^3^ arising from the unit conversion. Rearranging yields:

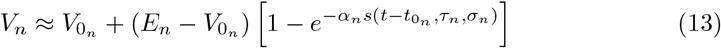

Since ionic conductance is modeled by the lognormal function, the corresponding state transition profile is represented by its cumulative distribution function (CDF):

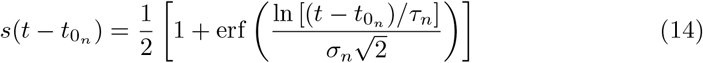

Equation (13) indicates that the sigmoid function *s*(*t*; …) is dimensionless and varies between 0 and 1. It describes the state transition of the membrane potential from its initial value 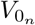 to its final value 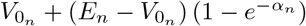.

The following model is proposed to describe the succession of these four state transitions:

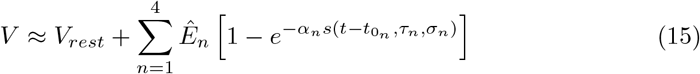

With 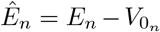 and 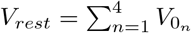 Ideally, the AP returns to its equilibrium level, so the sum of the magnitudes of the four successive state transitions equals zero.

An alternative formulation is obtained by defining *Êα*_*n*_ = *α*_*n*_*Ên* and applying an affine approximation of the exponential function:

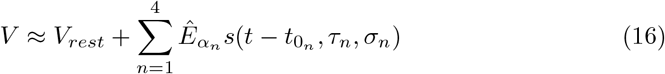

#### 2.3.1 Nernst Potential Estimation

The membrane potential *V* (*t*) is decomposed into four overlapping phases, each describing the dominant state transition driven by the conductance of a given ion. The membrane is modeled as a capacitor in parallel with four branches: three representing the ionic pathways of sodium, potassium, and leakage currents, and a fourth representing the sum of external currents responsible for depolarization. At rest, the three membrane branches are open. When *V* reaches the threshold associated with a given ion, its conductance becomes nonzero and the corresponding branch closes, allowing ionic passage. The branch reopens when *V* reaches the ion’s Nernst potential, measured relative to a zero-voltage reference. At rest, *V*_*rest*_ ≈ −70 mV. Depending on the direction of the current, the Nernst potential *E*_*n*_ is given by:

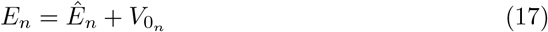

It is calculated relative to the value of *V* at the moment the conductance first becomes nonzero, i.e., at time 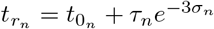 , such that 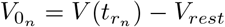

#### 2.3.2 Experimental Considerations

The Hodgkin-Huxley model was originally developed to describe ionic currents under voltage-clamp conditions, establishing the relationship between membrane conductances and action potential generation through a system of first-order differential equations. The present framework complements this approach by extending the analysis of neuronal bioelectrical activity toward quantitative processing of in vitro recordings. Equation (9) shows that the AP emerges from the superposition of three temporally shifted ionic fluxes, each representing a state transition of the membrane potential induced by a specific ionic species. Including the external ionic flux responsible for triggering depolarization, the AP waveform is described as a sequence of four successive state transitions — a characteristic feature of biological systems organized as networks of coupled stochastic processes whose collective behavior produces a stereotypical, deterministic free response. This model is applied to electrophysiological recordings from lamprey reticulospinal neurons following sensory stimulations that elicited limb movements.

#### 2.3.3 Experimental Setup

This study used AP recordings from a previous in vitro study [12], obtained from lamprey reticulospinal (RS) neurons depolarized by sensory stimulation (mechanical pressure on the head skin, electrical shocks, etc.) sustained for up to several minutes. Experiments were performed on a semi-intact preparation; recorded RS cells were located in the middle rhombencephalic reticular nuclei. Single and paired intracellular recordings were performed on the giant Müller cells using an Axoclamp 2A amplifier, Digidata 1322 interface, and Clampex 9.2 software. AP waveforms were extracted (Fig. 5) by segmenting the recordings into 12 ms windows centered on the AP peak, yielding 1413 spikes. Each spike was fitted with a signal-to-noise ratio SNR ≥ 40 dB (Fig. 6), defined as the logarithmic ratio of the energy of the analytical signal *s*_*a*_(*t*) to that of the residual error *s*_*a*_(*t*) − *s*_*n*_(*t*).

**Fig. 5:**
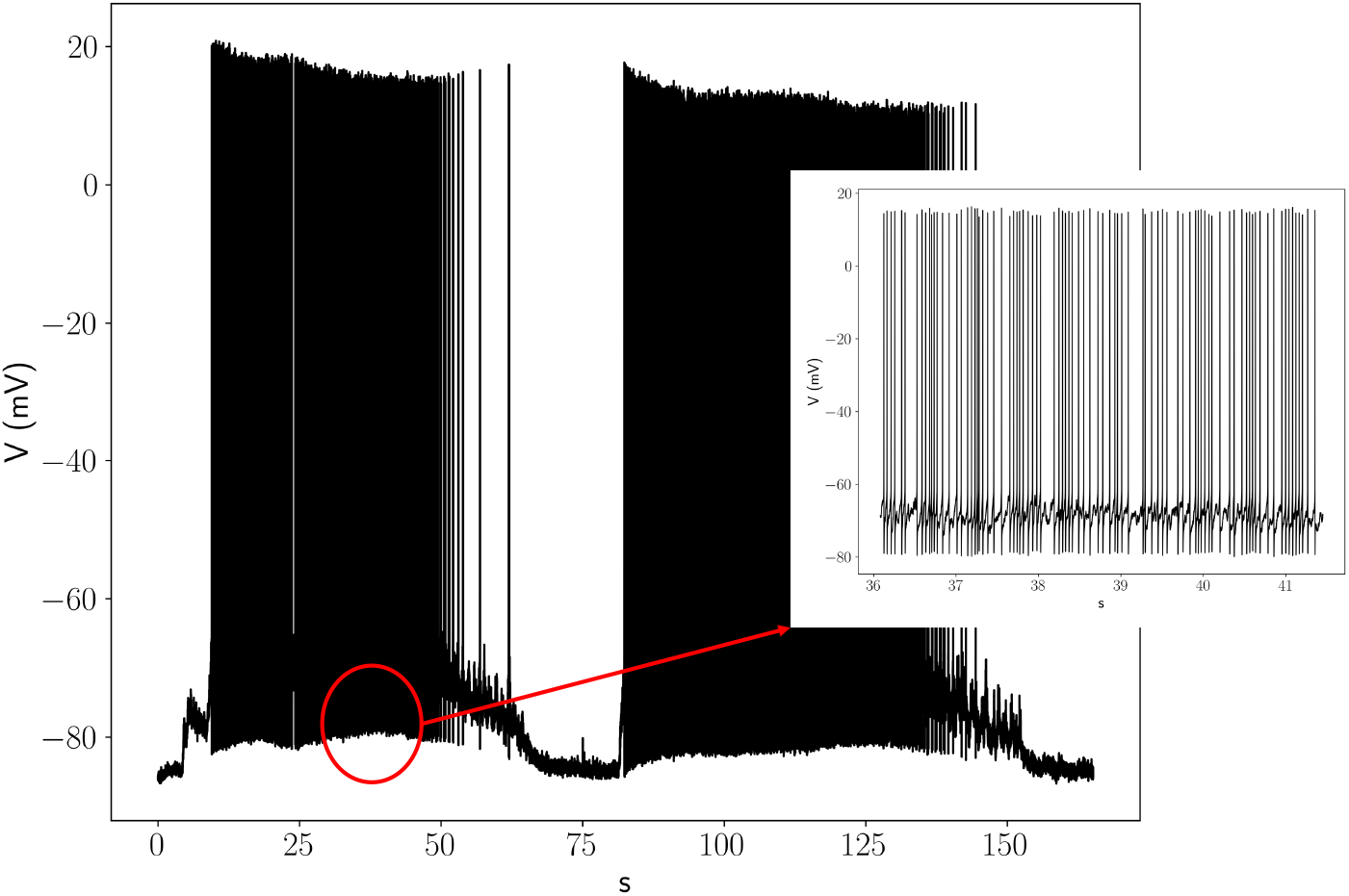
Experimental data used in this study: electrophysiological recordings of lamprey reticulospinal (RS) neurons following external stimulation [12], with an enlargement showing the AP train structure of the biosignal.

**Fig. 6:**
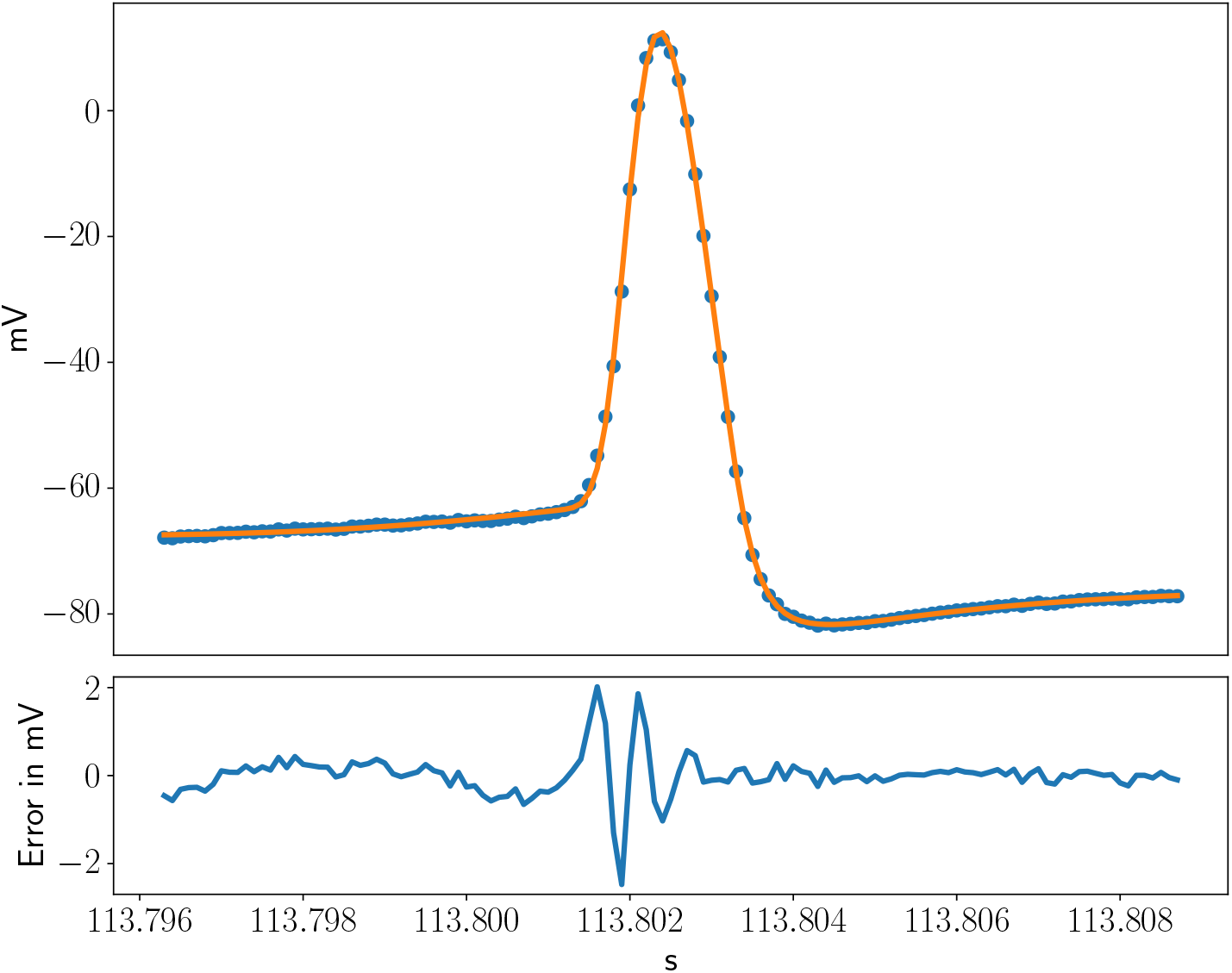
Representative fitting result of the proposed model applied to one of the 1413 action potentials in Fig. 5, with an SNR of 43.5 dB.

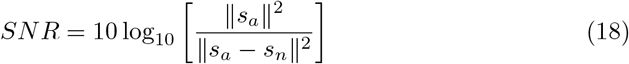

#### 2.3.4 Analysis of the Fitting Results

Each fit describes the AP profile using 21 parameters, as given by equation (15), 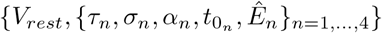 .These parameters quantify both the processes involved in neuronal depolarization and those underlying spike formation. The data come from lamprey RS cells stimulated by sensory inputs; the resulting spike train (Fig. 5) exhibits an inter-spike interval distribution (Fig. 14) confirming that AP generation is itself a stochastic process.

A representative fitting result is shown in Fig. 6, illustrating that the stereotypical AP waveform arises from three coupled stochastic processes governing the depolarization, repolarization, and return-to-rest transitions. The natural waveform variability reflects the intrinsic variability of the model parameters: biological variability in AP generation is thus embedded in the parameter distributions. Fig. 7 superimposes all recorded APs with their mean profile, computed from average parameter values, while Fig. 8 depicts the average profiles of the four state transitions.

**Fig. 7:**
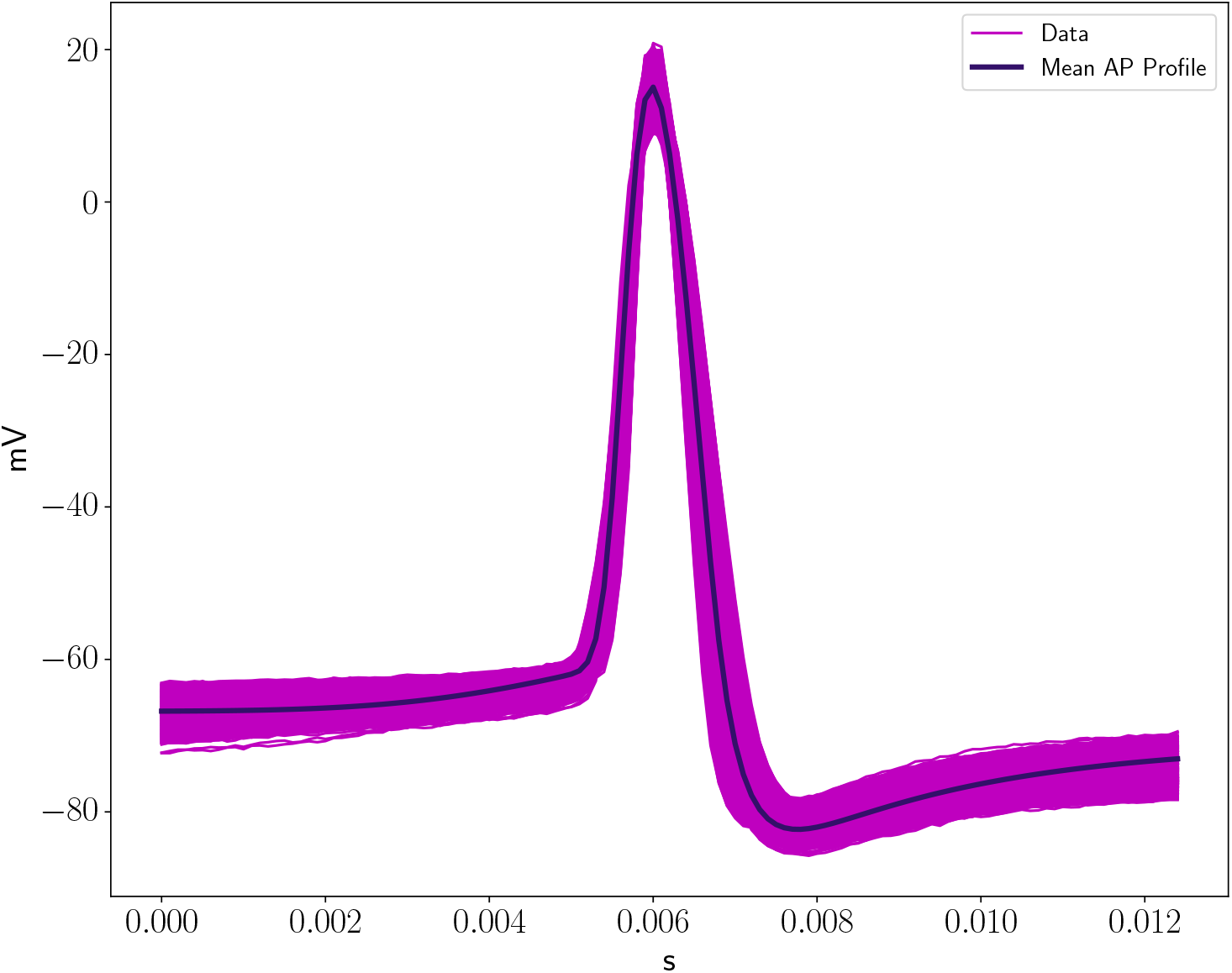
Superposition of the 1413 AP waveforms with the mean profile computed from the average values of the 21 model parameters.

**Fig. 8:**
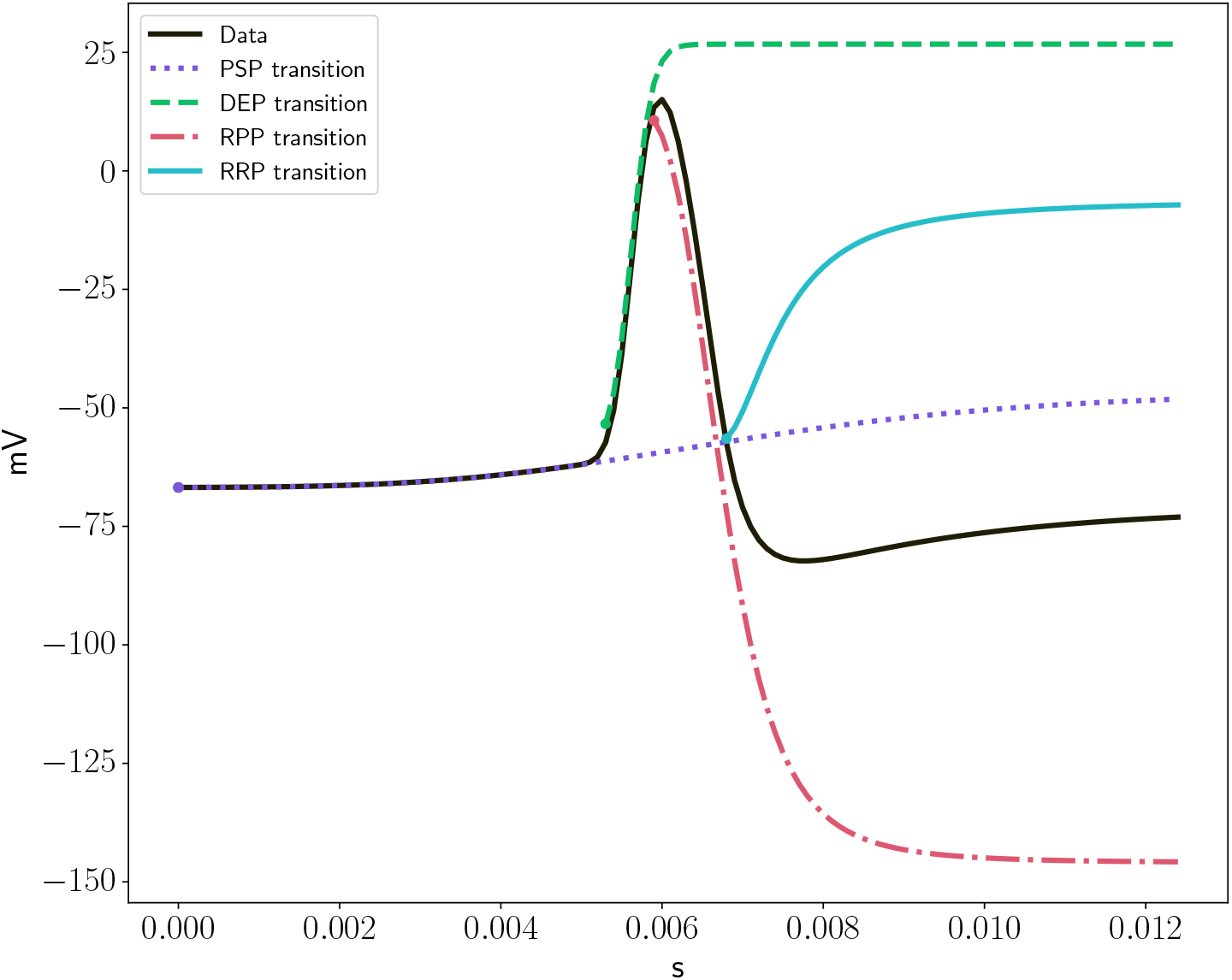
Mean AP waveform and its four constituent state transitions (PSP, DEP, RCP, RRP) as described by the proposed model.

From the mean parameter values, the proposed model yields the dominant sodium and potassium conductances (Fig. 9), as well as the distributions of the Nernst potentials and net ionic charges during depolarization and repolarization (Figs. 10–13). Since these distributions are associated with causal, ergodic biological systems and are typically positively skewed, each parameter *p* is fitted with a lognormal function 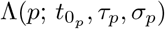 , from which the mean 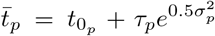and standard deviation 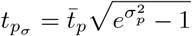 are derived directly.

**Fig. 9:**
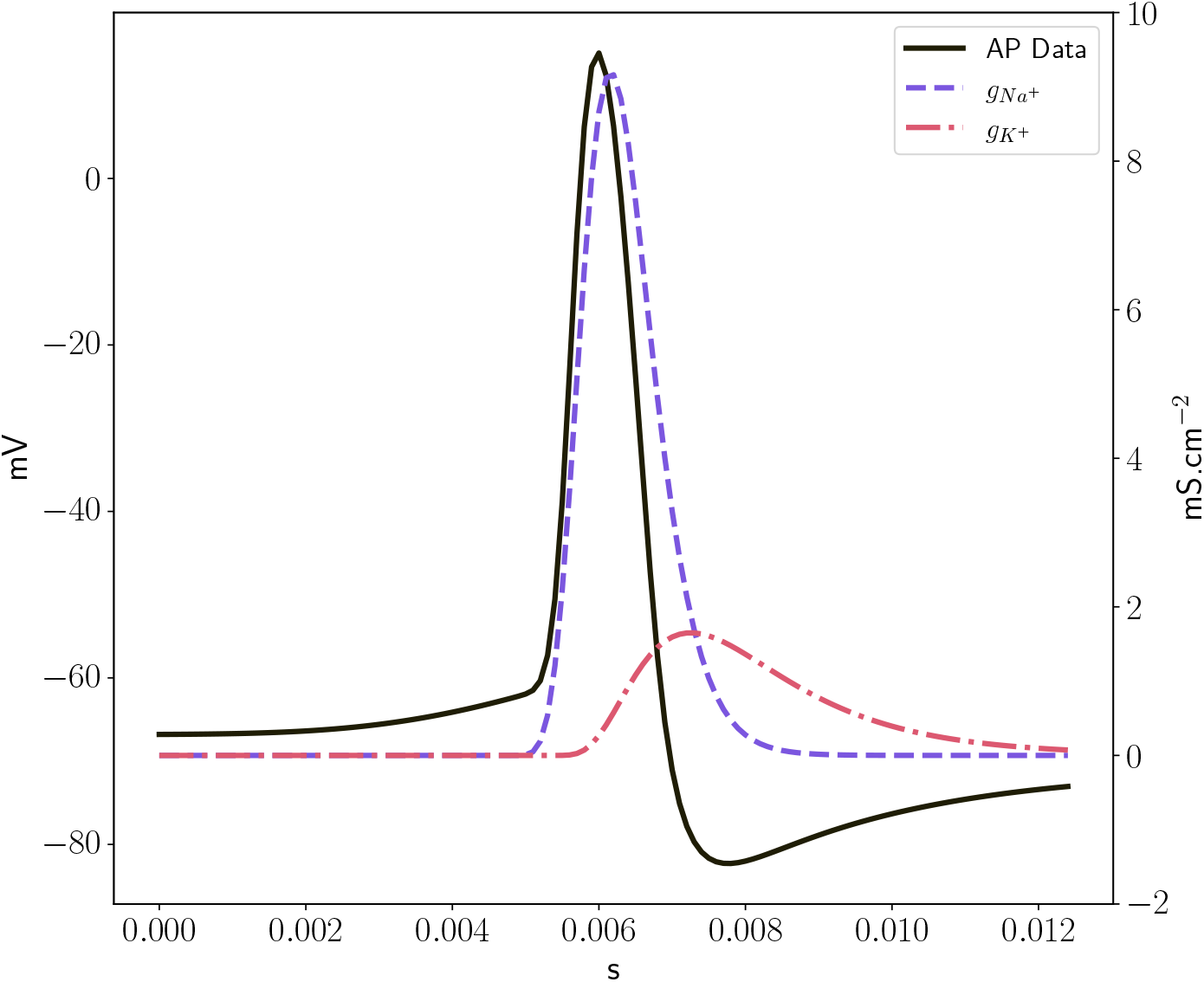
Sodium and potassium conductance waveforms associated with the AP, computed from the lognormal model parameters.

**Fig. 10:**
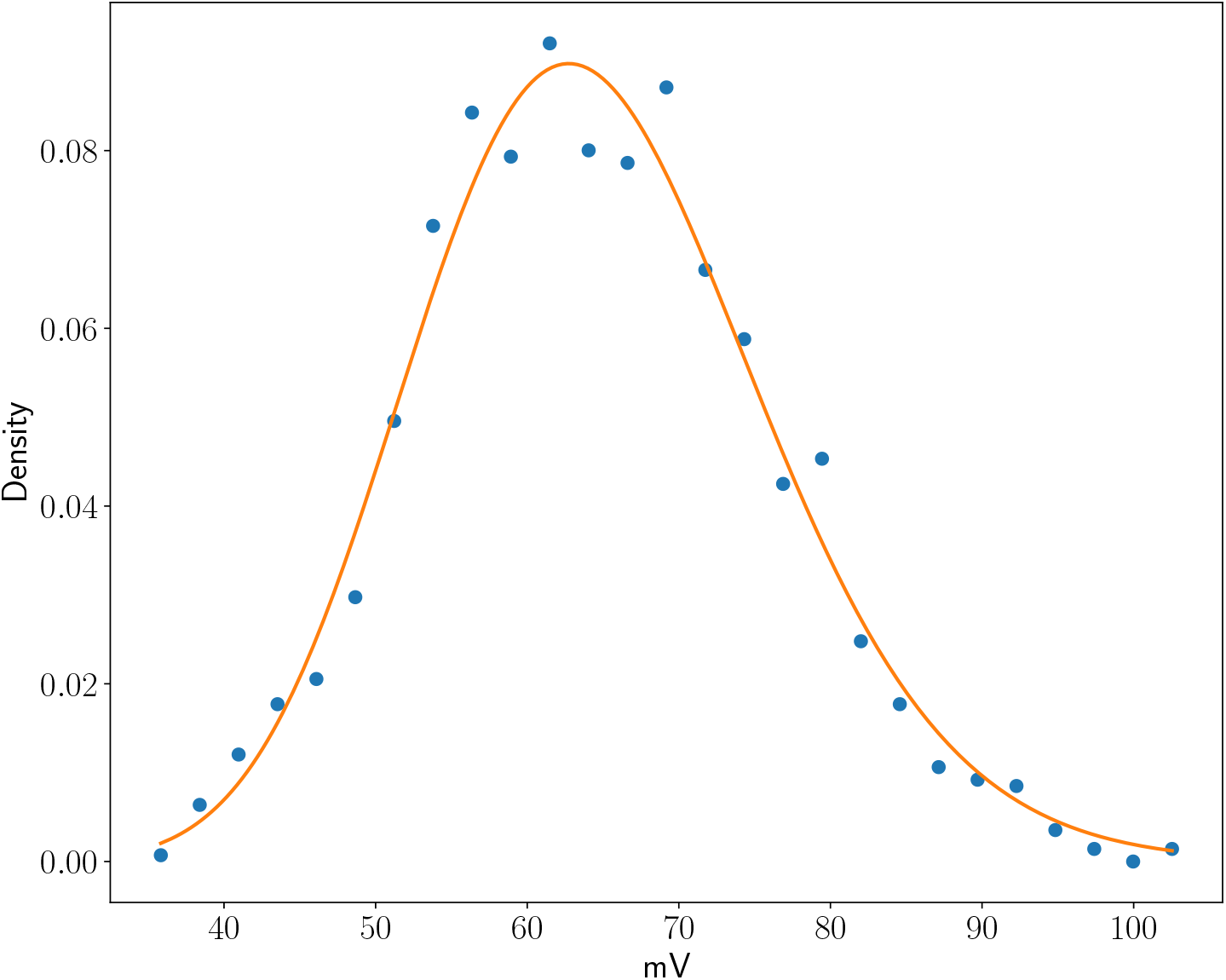
Distribution of the Nernst potential associated with the depolarization phase (DEP), corresponding to Na^+^, with mean value 65.4 mV (Table 1).

**Fig. 11:**
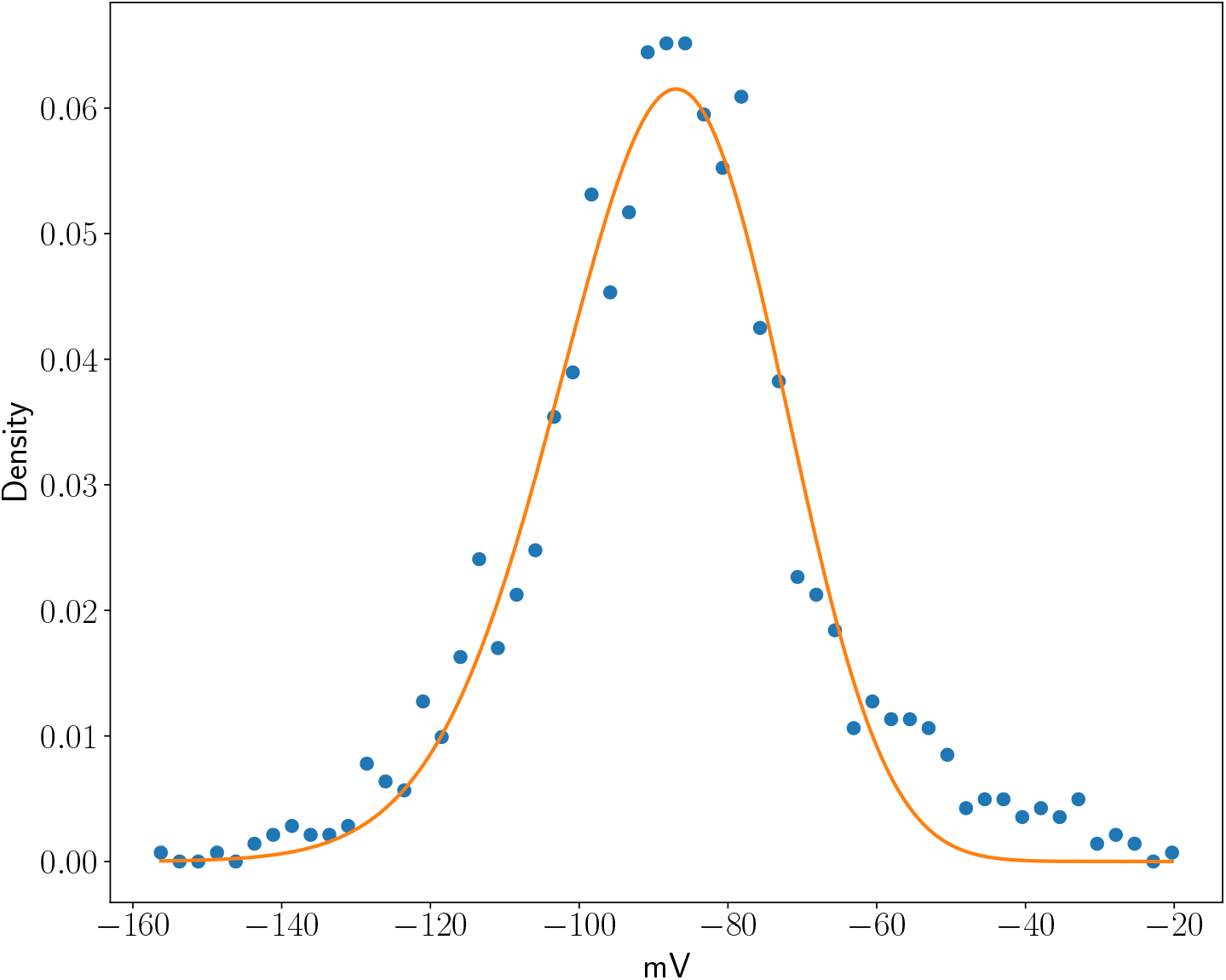
Distribution of the Nernst potential associated with the repolarization phase (RPP), corresponding to K^+^, with mean value −89.8 mV (Table 1).

**Fig. 12:**
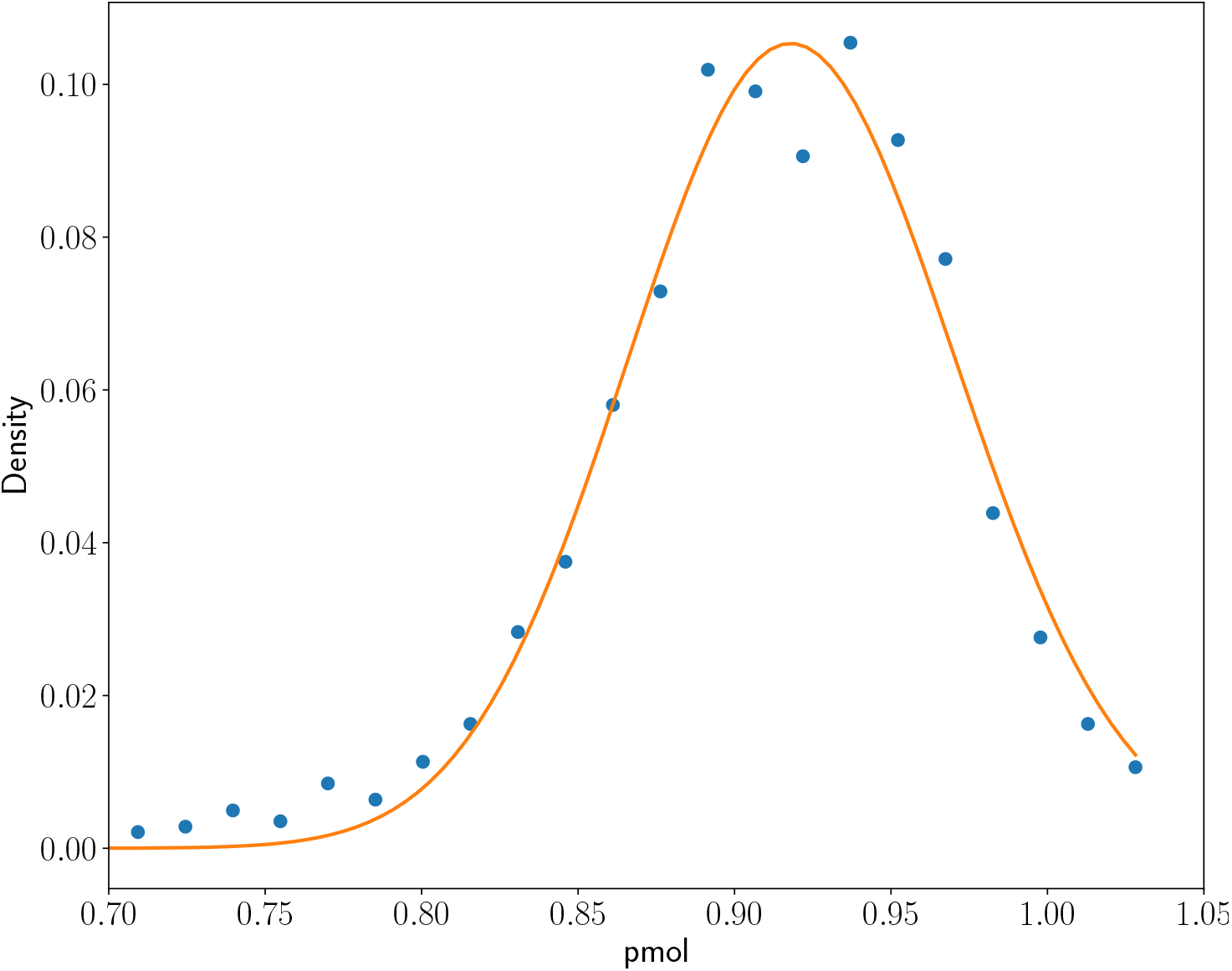
Distribution of the net Na^+^ charge involved in the depolarization phase, with mean value 0.92 pmol (Table 1).

**Fig. 13:**
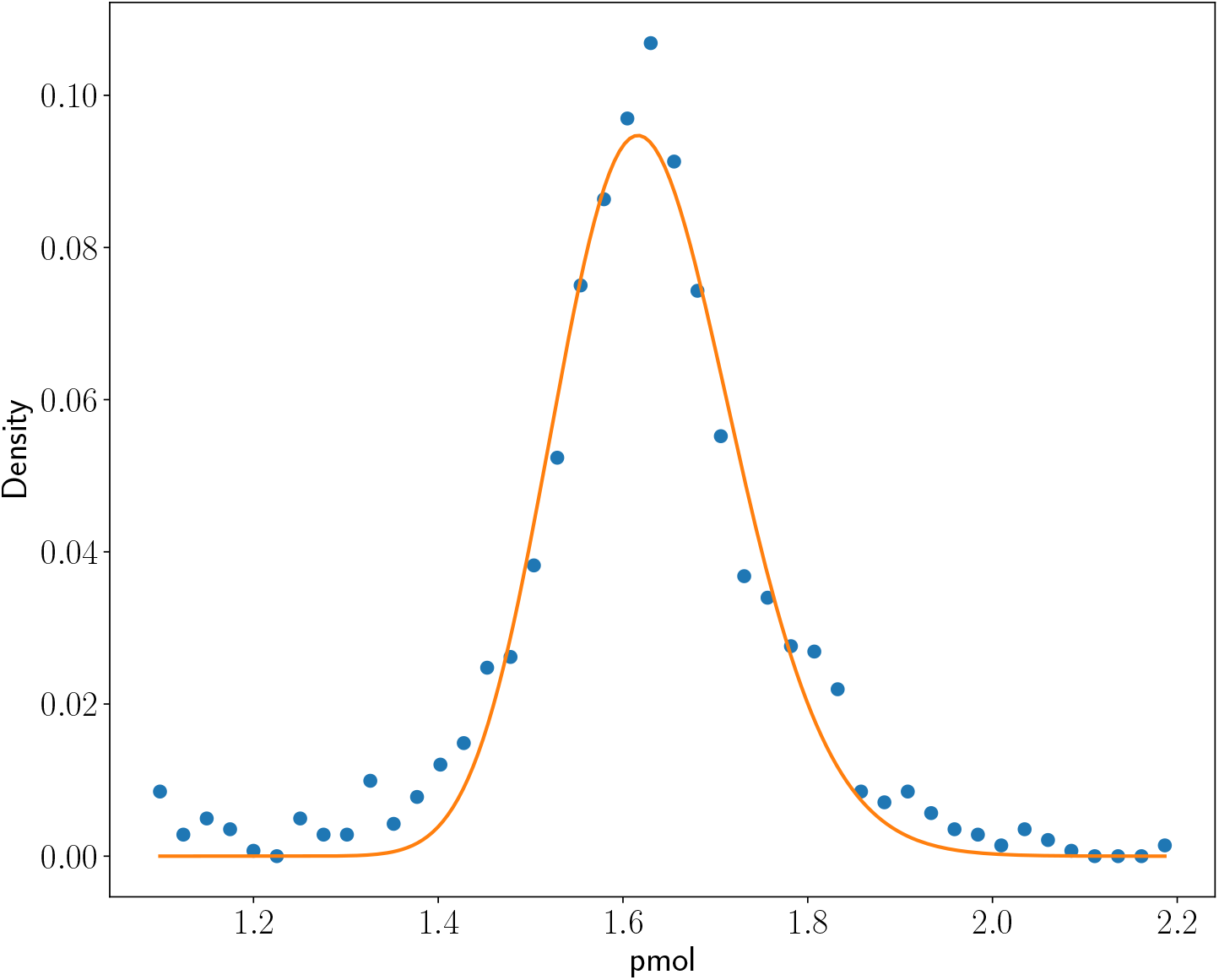
Distribution of the net K^+^ charge involved in the repolarization phase, with mean value 1.63 pmol (Table 1).

**Fig. 14:**
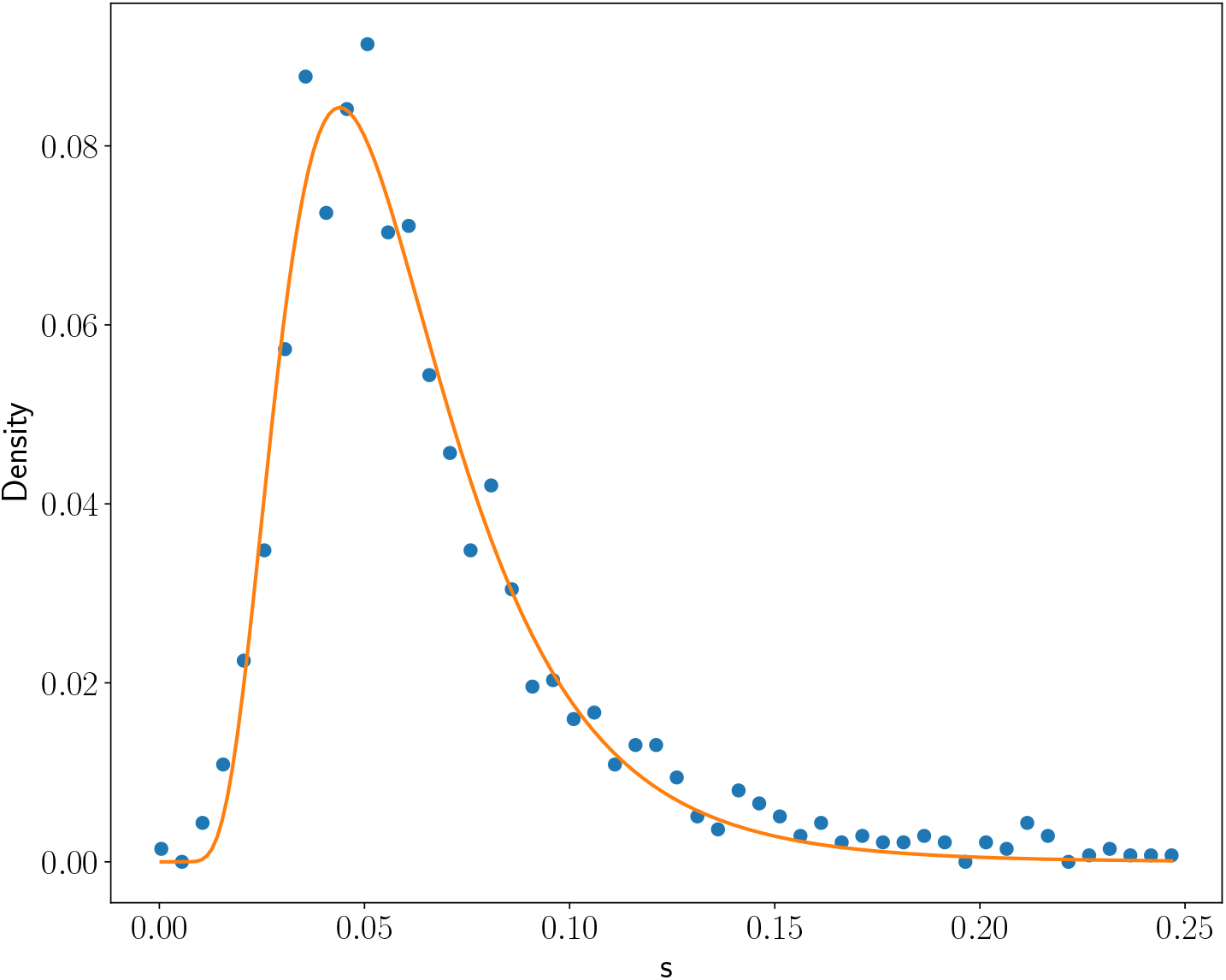
Distribution of inter-spike intervals associated with the AP depolarization onset time *t*_0_ following external stimulation. The mean inter-spike interval is 69.3 ms, corresponding to an average firing rate of 14.4 Hz.

It is important to distinguish between the lognormal modeling of conductance — which describes the temporal random variable governing ionic transmembrane transit — and the lognormal modeling of parameter distributions, which characterizes the biological variability associated with AP generation.

#### 2.3.5 Depolarization Markers

The estimated parameters are used to construct the following markers, characterizing the three main state transitions of the membrane potential underlying AP formation. Each transition of order *n* is described by:

1. State Transition Onset Time: 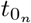
2. Time of Peak Conductance Rate: 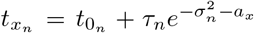 with 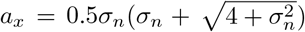
3. Effective Conductance Duration: *T*_*n*_ = 2*τ*_*n*_ sinh 3*σ*_*n*_
4. Post-Trigger Conductance Inertia Time: 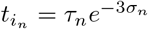
5. Conductance Reaction Time: 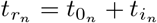
6. Nernst potential: 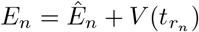
7. State Transition Amplitude: 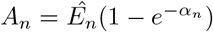
8. Net Ionic Charge Associated with the State Transition (pmol): *Q*_*n*_ = 0.01036*A*_*n*_

The effective depolarization threshold is estimated as *V*_*th*_ = *V* (*t*_*th*_) − *V*_*rest*_, where 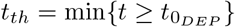 is determined from the onset time of the DEP state transition.

Like the HH model, the proposed model exhibits parameter degeneracy: multiple parameter combinations can yield indistinguishable AP waveforms. For instance, infinitely many combinations of *t*_0_, *τ* , and *σ* produce the same conductance peak time 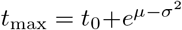 . In the absence of prior knowledge constraining parameter ranges, the decomposition of AP recordings into sigmoid-like components was guided by known physiological quantities, such as the Nernst potentials of sodium and potassium and the approximate resting potential. Decomposing the 1,417 AP profiles into four membrane potential state transitions yielded the mean parameter and marker values from their estimated distributions.

Table 1 summarizes the average marker values, providing insight into the stochastic processes underlying AP generation and their coupling to produce a stereotypical waveform. Note that the onset time *t*_0_ denotes the instant at which a state transition is triggered, whereas the reaction time *t*_*r*_ marks the instant at which it becomes observable; the interval between them defines the inertia latency *t*_*i*_ of the transition. The following observations are drawn from the estimated parameter values.

1. The estimated Nernst potential for sodium, *E*_*Na*_ ≈ 65.4 mV, is consistent with the Nernst equation evaluated at 10°C, as typically used in this type of in vitro experiment.
2. During the repolarization phase (RPP), dominated by K^+^ efflux, the estimated potential *E*_*K*_ ≈ −89.8 mV is in close agreement with the theoretical K^+^ Nernst potential of −90 mV.
3. Depolarization is triggered when the membrane potential exceeds *V*_*rest*_ by approximately *V*_*th*_ = 9 mV.
4. Repolarization onset coincides with the effective start of depolarization 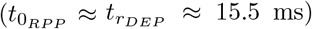, and repolarization becomes observable when depolarization peaks 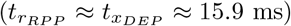 The return to rest is triggered near the peak of repolarization 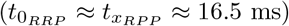.
5. The repolarization phase lasts approximately three times longer than depolarization and involves a greater net ionic charge transfer.
6. The ratio *η* = *T/t*_*i*_ differs markedly between phases: 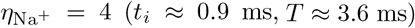 versus 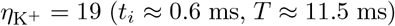, reflecting a larger sodium inertia relative to its shorter active duration, contrasting with a smaller potassium inertia but much longer conductance persistence.
7. The net balance of all ionic fluxes contributing to AP generation is zero to within 0.1 pmol (Table 1).
8. The four state transitions follow the sequence: 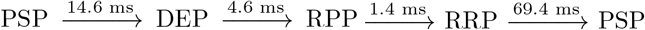
9. The mean interspike interval is 69.4 ms, corresponding to an average firing rate of 14.4 Hz.

### 2.4 Analysis of State Transition Coupling

The AP waveform emerges from a precise coupling among four successive state transitions: the onset of repolarization (RPP) coincides with the reaction time of depolarization (DEP), while the first inflection point of repolarization triggers the return-to-rest transition (RRP).

Analysis of the onset-time distributions for the DEP, RPP, and RRP phases reveals progressively shifted and broadened distributions, with mean values of 14.6, 15.4, and 16.8 ms and standard deviations of 7.16, 7.24, and 7.28 ms, respectively. The coefficients of variation (0.49, 0.47, and 0.43) are all well below unity, indicating that the inter-phase coupling is stochastic yet unimodal.

Based on the shapes and relative positions of these distributions, the coupling between successive transitions is modeled as a convolution product (Fig. 15). As illustrated in Fig. 16, each depolarizing event sequentially triggers a set of interconnected stochastic processes. This triggering event is represented symbolically by a Dirac delta function *δ*(*t*−*t*_0_); since the event is stochastic in nature, the delta function is more appropriately interpreted as a distribution with mean *t*_0_.

**Fig. 15:**
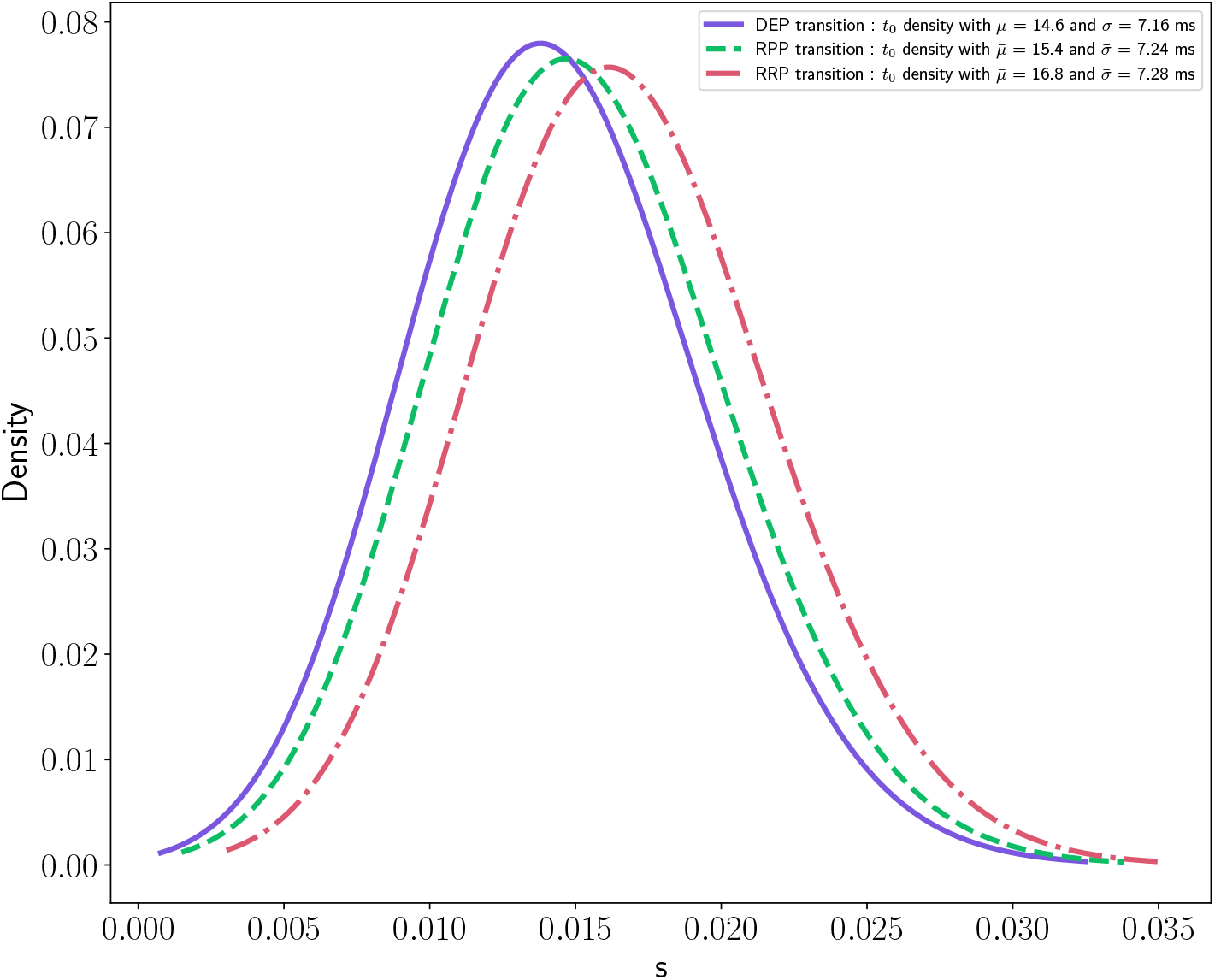
Distributions of the onset times *t*_0_ of the three AP state transitions: depolarization (DEP), repolarization (RPP), and return to rest (RRP), characterizing the stochastic processes governing their sequential coupling.

**Fig. 16:**
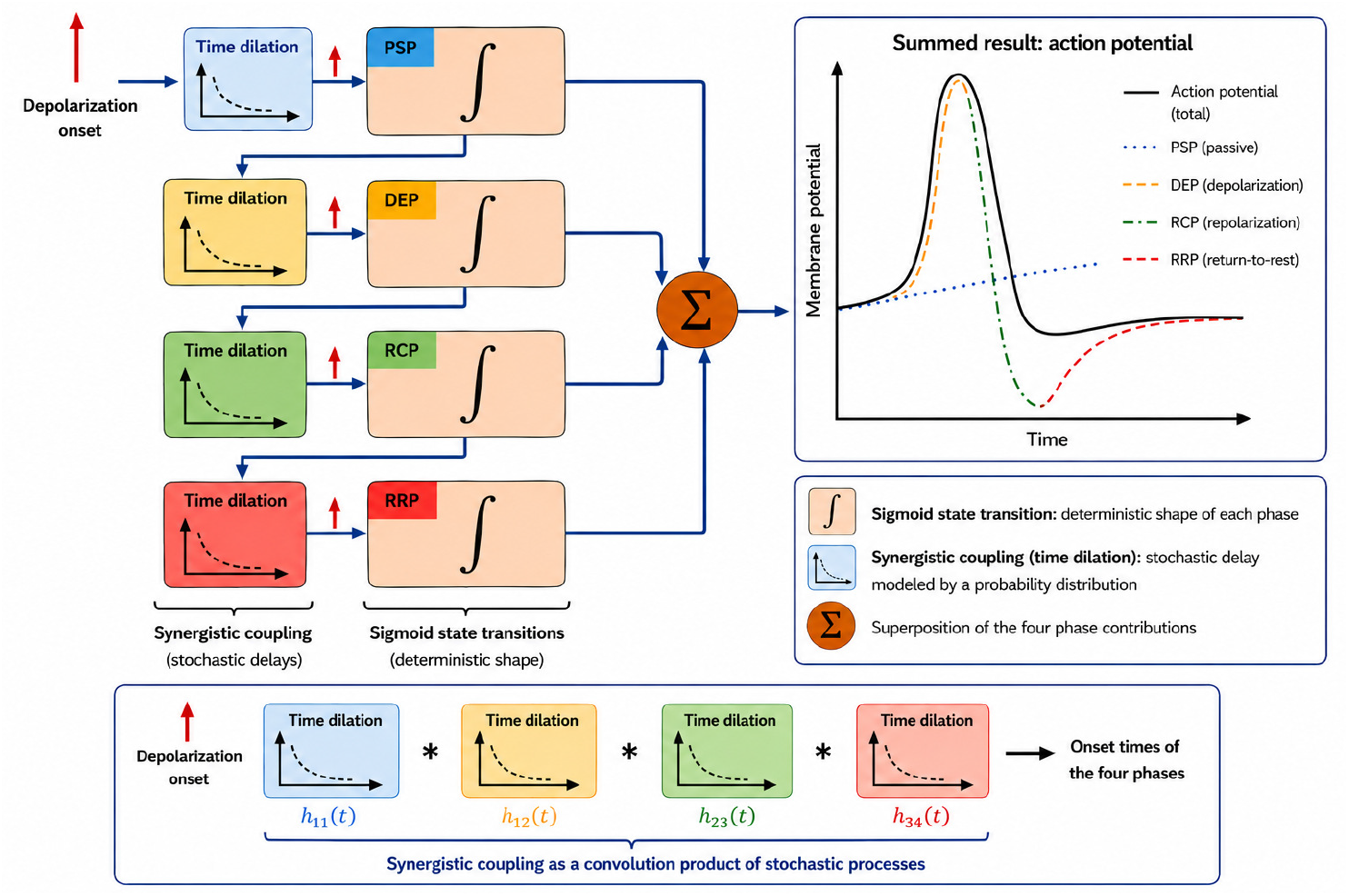
Convolution-based representation of the stochastic processes governing the sequential coupling among AP state transitions associated with different ionic conductances.

The depolarizing event first triggers the stochastic process governing sodium channel opening — an intermediate control layer between membrane potential and channel state. Repeated AP generation enables the construction, through superposition, of onset-time distributions for each phase. When these distributions are convolved, both their means and dispersions increase sequentially.

Formally, let *h*_*i*_ and *h*_*j*_ denote the onset-time distributions of two coupled state transitions, with means 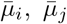 and variances 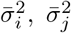. Their coupling is modeled by a stochastic delay process *h*_*ij*_ such that *h*_*j*_ = *h*_*i*_ ∗ *h*_*ij*_, with mean 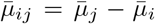and variance 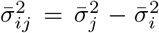. From the results in Fig. 15, the coupling distributions *h*_*ij*_ have mean values of 0.8 and 1.4 ms with standard deviations of approximately 1.1 and 0.8 ms, respectively. The coefficient of variation of 1.38 associated with the RCP → RRP coupling suggests that this process may represent a mixture of underlying stochastic processes.

In summary, the main three-state model not only characterizes the stochastic processes governing each AP phase, but also reveals that the couplings between successive transitions are themselves stochastic processes, embedded in the membrane’s interconnected biological structure.

### 2.5 Self-Similarity in Neuronal Organization

Analysis of the HH model reveals a hierarchical and parallel organization of ion channels, each comprising a network of highly interconnected gates whose conduction pathway is stochastic. Ionic conduction thus manifests as a spike train with interspike intervals governed by a lognormal distribution. An analogous phenomenon is observed at the neuronal scale: AP trains likewise follow a lognormal probability law. Just as membrane depolarization gives rise to ionic conductance, a neural flux triggered by stimulation corresponds to a state transition induced by internal or external stimuli.

At a higher scale, neuronal networks are themselves highly interconnected, with each neuron receiving inputs from, and projecting to, multiple others. In neuromotor networks, for instance, the velocity profiles of rapid ballistic movements follow a lognormal distribution [13], and the execution of such movements can be interpreted as a state transition induced by the depolarization of a specific neuromotor network configuration [17].

From a morphological perspective, biological systems often exhibit fractal-like structures reflecting self-similarity across scales. The present results support the view that cross-scale transitions — within the framework of the central limit theorem — are governed by a principle of functional self-similarity, with the lognormal distribution emerging as the common primitive across organizational levels.

## Discussion

Building on the Hodgkin-Huxley (HH) models of ionic conductance and action potential, an updated framework is proposed. It rests on the hypothesis that the bioelectrical functioning of the neuron results from strongly coupled stochastic processes, whose deterministic behavior emerges from the direct realization of their governing probability distributions, where the random variable represents the transmembrane transit time of ions. The sodium and potassium conductance profiles initially proposed by HH exhibit a probability density profile converging toward a lognormal limiting distribution, as confirmed by the fitting results shown in Figs. 1 and 2. The estimated model parameters quantify the relationship between ionic conductance and membrane voltage. In voltage-clamp experiments [4], sodium conductance data reveal a minimum depolarization threshold of 25 mV: above this value, conductance amplitude increases while its duration and response time decrease sharply, with onset time remaining constant relative to depolarization onset. Beyond 75 mV, conductance amplitude abruptly decreases, confirming that high depolarization levels progressively inhibit sodium conductance (Fig. 3).

The original HH action potential (AP) equation was used to construct an analytical model representing the AP waveform as four successive state transitions of the membrane potential *V* . The first transition represents the external stochastic process triggering neuronal depolarization; the remaining three describe, in succession, the depolarization, repolarization, and return-to-rest phases. Each phase corresponds to a transition of the equivalent membrane capacitor voltage between two equilibrium states. Because the membrane does not respond instantaneously, each transition undergoes three temporal dilations. The first is a pure delay *t*_0_, representing the interval between the onset of the triggering event and that of the corresponding transition. The second is the inertia time *t*_*i*_, characterizing the latency required for ionic channels to open and initiate conduction. The third corresponds to the conductance process itself, which transforms the ideal step-like voltage transition into a sigmoid-shaped temporal evolution. As described by equation (14), the AP results from the concatenation, with partial overlap, of these four state transitions, each modeled by five parameters (*τ, σ, α, t*_0_, *E*_*n*_), from which several physical quantities associated with AP initiation and formation can be estimated.

The proposed model enables direct analysis of an activated neuron’s bioelectrical properties from electrophysiological recordings. Applied to recordings from lamprey reticulospinal (RS) neurons following external sensory stimulation, it allows the indirect estimation of physiologically meaningful quantities, including the Nernst potentials of major ionic species, the mean membrane depolarization threshold, the resting membrane potential, and the net ionic charges contributing to AP generation. These results open new perspectives for artificially generating desired bioelectrical responses through controlled manipulation of model parameters. More broadly, the state-transition framework can be readily extended to other biosignal classes, including electrocardiographic (ECG) recordings.

The lognormal profile of ionic conductance motivates interpreting the primitive underlying biosignals as the direct realization of a cumulative distribution function of a temporal random variable associated with one or more stochastic processes. Such realization emerges from the highly interconnected, hierarchical, and parallel organization of the constituent elements, whereby stochastic microscopic activity gives rise to deterministic collective behavior. A second key observation from the HH study is the pervasive presence of agonist-antagonist couplings between processes: the activity of one process triggers a response in another, ultimately restoring the system’s equilibrium. In rhythmic biological systems, multiple coupled stochastic processes ensure a deterministic functional outcome. The biosignal corresponds to the repeated expression of a complex pattern representable as a sequence of state transitions, exhibiting variability around a dominant operating point — as seen in cardiac rhythm maintenance or in AP trains driving limb movements through muscle contractions.

This reveals how biological systems operate stochastically at the microscopic level yet exhibit deterministic, predictable behavior at higher organizational scales, with variability confined within the normal operating range. Similar biological entities self-organize and operate in parallel to collectively realize the cumulative distribution function of their governing probability law, applying this output to a terminal cumulative effector that integrates a large number of infinitesimal contributions to produce a deterministic state transition. An illustrative example is provided in [14], where AP trains generated by lamprey RS neurons are transmitted to muscle fibers, producing limb movements in response to sensory stimulation.

The lognormal function is therefore selected as the fundamental primitive of biosignals. Depending on the nature of the signal, it may manifest either as a probability density function or as a cumulative distribution function. In the HH study, conductance data recorded via ionic currents provide direct access to the density profile, whereas AP data obtained by measuring the voltage across the membrane capacitor *C*_*m*_ provide access to its cumulative distribution function. Within neuroscience, Buzsáki and Mizuseki [15] reviewed the lognormal distribution as a descriptor of anatomical and physiological quantities in neuronal systems. More recently, [16] demonstrated that the sinusoid, commonly used in spectral decomposition of biosignals, can be interpreted as the limit of an infinite concatenation of Gaussian densities — and, accounting for the causal nature of biological responses, more faithfully as an infinite concatenation of lognormal densities. The sinusoid and its variants (wavelets, etc.) thus appear to be composed of a more elementary primitive arising from stochastic biological activity. For signals that are not only rhythmic but also transient and of finite energy, the lognormal function constitutes a more appropriate primitive.

As a perspective for neuroscience, the AP constitutes the fundamental unit of information transmission in neuronal networks. APs occur at random times, encoding information in inter-spike intervals that depend on synaptic connectivity, depolarization threshold, and other physiological parameters. The proposed model can generate spike trains whose inter-spike intervals follow specific probability laws, controlled through the depolarization markers described above. This enables simulation of signal propagation through a network with prescribed depolarization thresholds, synchronization delays, or ionic quantities confined to defined operating ranges — reproducing neuronal network behavior across physiological parameter variations and generating biosignal patterns associated with experimentally observed pathological conditions. Figure 17 illustrates an artificially generated AP train produced using the proposed model, with parameters randomly selected within their estimated variability ranges.

**Fig. 17:**
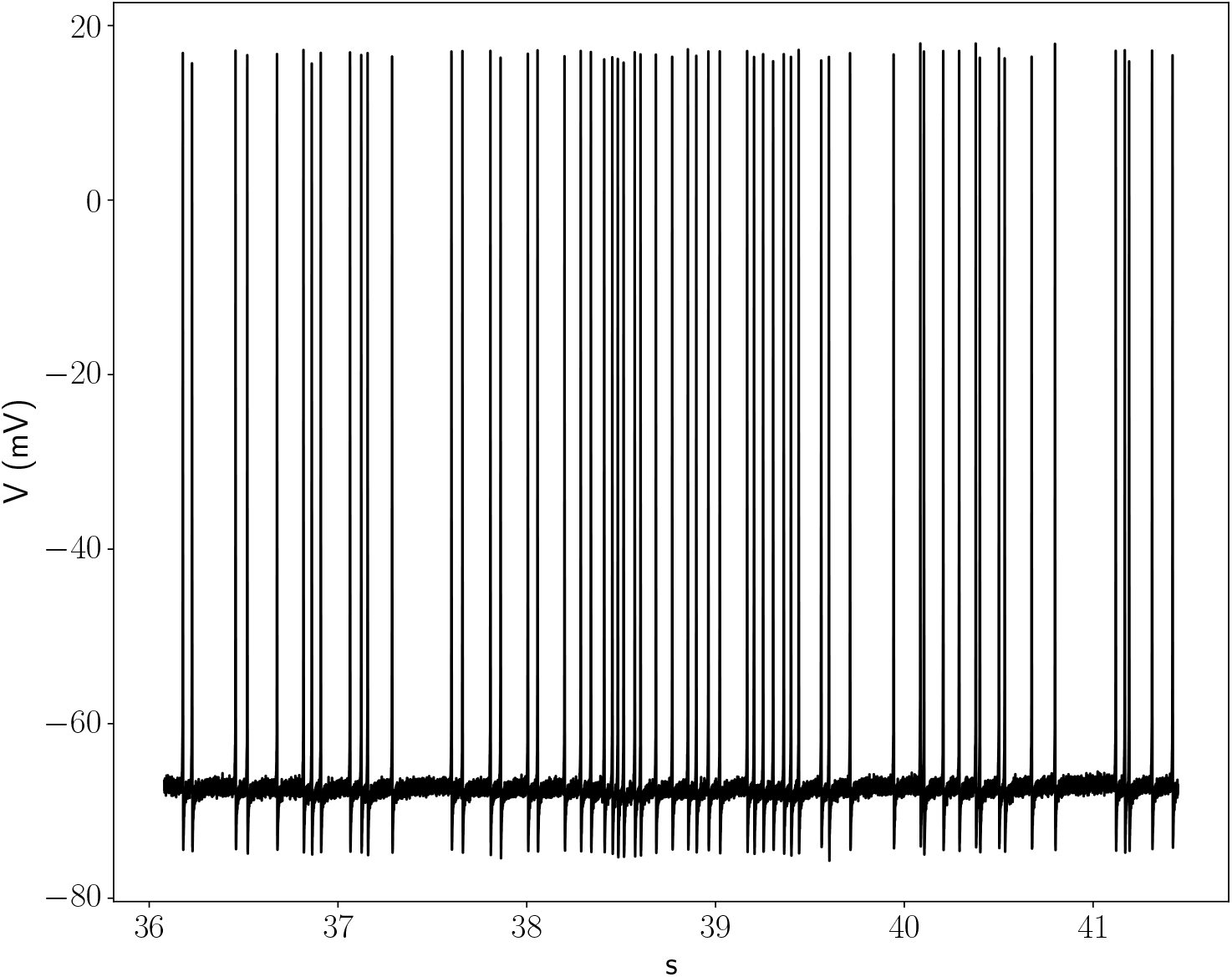
Synthetic reconstruction of lamprey reticulospinal (RS) neuron activity generated by the proposed model, reproducing a signal comparable to that in Fig. 5.

## 4 Acknowledgements

AI-assisted tools were used for language refinement of the manuscript and graphic illustration. The author takes full responsibility for the scientific content.

